# DECONbench: a benchmarking platform dedicated to deconvolution methods for tumor heterogeneity quantification

**DOI:** 10.1101/2020.06.06.131482

**Authors:** Clémentine Decamps, Alexis Arnaud, Florent Petitprez, Mira Ayadi, Aurélia Baurès, Lucile Armenoult, HADACA consortium, Rémy Nicolle, Richard Tomasini, Aurélien de Reyniès, Jérôme Cros, Yuna Blum, Magali Richard

## Abstract

**Motivation:** Quantification of tumor heterogeneity is essential to better understand cancer progressionand to adapt therapeutic treatments to patient specificities.

**Results:** We present DECONbench, a web-based application to benchmark computational methods dedicated to quantify of cell-type heterogeneity in cancer. DECONbench includes benchmark datasets, computational methods and performance evaluation. It allows submission of new methods.

**Availability and implementation:** DECONbench is hosted on the open source codalab competition platform. It is freely available at: https://competitions.codalab.org/competitions/23660.

**Supplementary information:** Additional information is available online and on our website: https://cancer-heterogeneity.github.io/deconbench.html.

## 1 Introduction

Since the recent development of high-throughput sequencing technologies, cancer research has focused on characterizing the genetic and epigenetic changes that contribute to the disease. However, these studies often neglect the fact that tumors are constituted of cells with different identities and origins. Quantification of tumor heterogeneity is of utmost interest to the bioinformatics and biomedical research community as multiple components of a tumor are key factors in tumor progression, clinical outcome and response to therapy. Advanced microdissection techniques to isolate a population of interest from heterogeneous clinical tissue samples are not feasible in daily practice. Single-cell technologies, while promising, have intensive protocols and require expensive and specialized resources, currently hindering their establishment in a clinical setting (Avila Cobos et al., 2018). An alternative is to rely on deconvolution methods that infer cell-type composition in silico. Bioinformatics tools to assess the different cell populations from bulk transcriptome (Becht et al., 2016; Nazarov et al., 2019; Blum et al., 2019) and methylome (Houseman et al., 2014; Lutsik et al., 2017; Decamps et al., 2020) samples have been recently developed, including reference-based and reference-free methods. This offers several advantages, notably the possibility to re-analyse a large number of publicly available datasets. However, their efficacy assessment has been impaired by the lack of dedicated benchmarking studies, which is often the case for methodological developments (Ellrott et al., 2019). Here we present DECONbench, an innovative public digital benchmarking platform, open source and freely available, aiming to compare deconvolution methods for tumor heterogeneity quantification. It includes benchmarking datasets, state-of-the-art computational methods and it enables the submission of new methods.

## 2 The benchmarking platform infrastructure

DECONbench takes advantage of the Codalab web-based platform (https://competitions.codalab.org/) to provide a common software environment for evaluating deconvolution methods. Users submit a full R program that is applied to the provided benchmark datasets and compared to the ground truth. DECONbench outputs a performance score displayed on the leaderboard (Fig. 1).

**Fig. 1.**
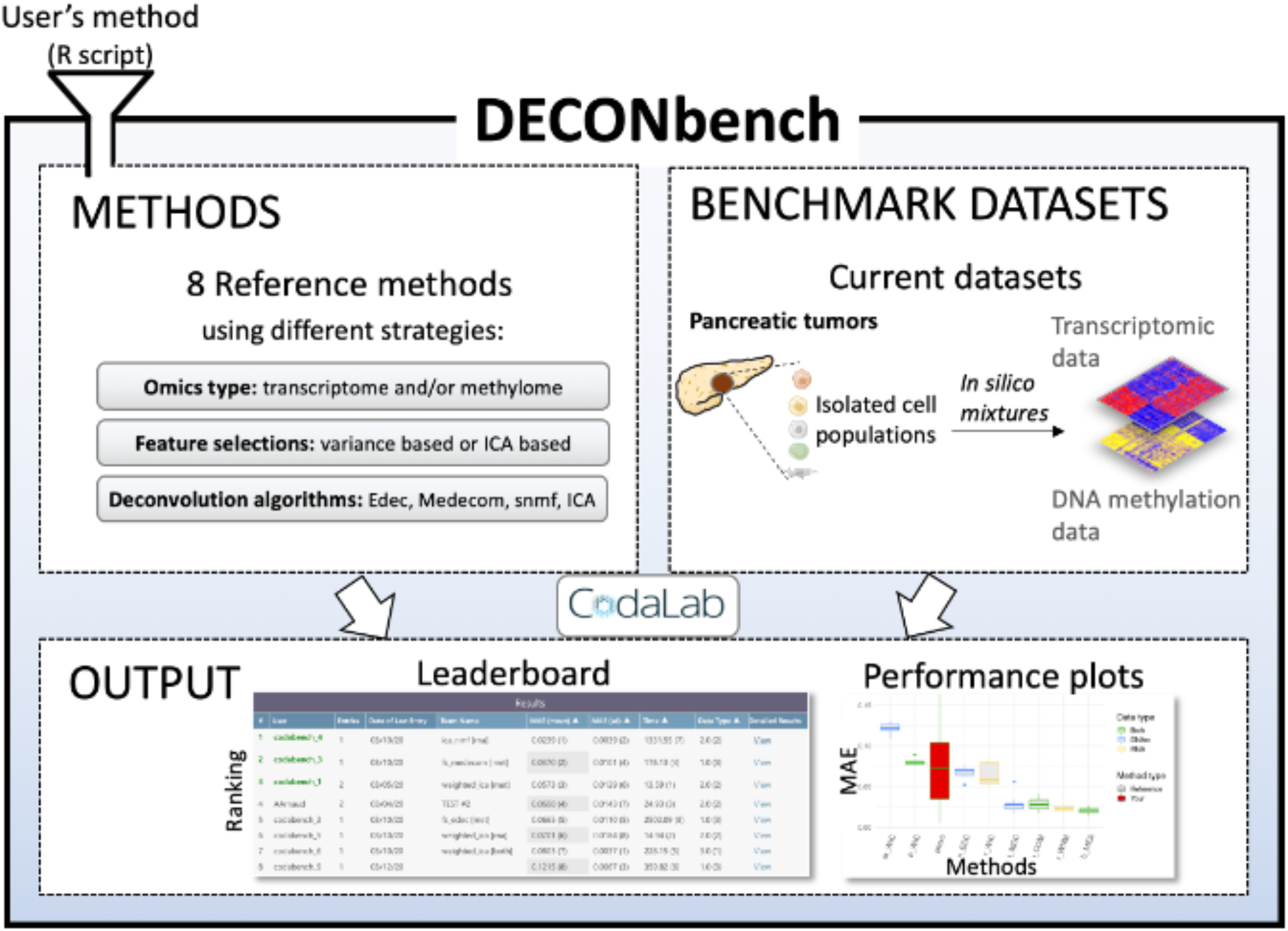
Overview of the DECONbench platform. The platform proposes a set of 8 reference deconvolution methods and benchmark datasets consisting of paired methylome and transcriptome of in silico mixtures from pancreatic tumors. The platform outputs the performance of each method on a leaderboard and provides plots for deeper evaluation. New methods are automatically compared to the existing ones.

## 3 Current benchmark datasets and methods

We have generated transcriptome and methylome benchmarking datasets from primary cells from pancreatic tumors and sorted cells from public datasets (supplementary Fig. 1). Heterogeneous samples were simulated using mixture of individual cell populations. Sample compositions are not accessible to the users. Methods are evaluated on their accuracy to estimate the cell-type proportion per sample from transcriptome and/or methylome heterogeneous profiles. The discriminating metric is the mean absolute error between the estimate and the ground truth. We recently used this unreleased dataset in a data challenge (https://tinyurl.com/hadaca2019). The best methods collectively discovered during the challenge are provided on DECONbench as reference methods (supplementary Table 1). The methods consist of various statistical approaches and novel strategies integrating transcriptome and methylome.

## 4 Usage

DECONbench is designed to execute methods developed in R statistical programming language, using a docker image provided on our website. A list of R packages installed on the docker image is as well provided. Users need to: i) register to DECONbench on the participate tab, download the starting kit and the public datasets ii) develop an algorithm according to DECONbench guidelines and iii) submit their code (a zip file) in the participate tab. Submitted algorithms are evaluated on DECONbench datasets and benchmarked with the other methods. Resulting scores appear on the leaderboard and a fact sheet is edited summarizing the performances (Supplementary Fig. 2). Importantly, users can choose whether they want their algorithm to be public or private.

## 5 Perspectives

This platform is a unique opportunity to compare the performance of deconvolution methods on different omics data. It can be used to assess the performance of newly developed methods by applying them on high quality benchmark datasets in a user-friendly fashion. The structure of DECONbench is open to evolution. Work is ongoing to generate new benchmark datasets that will be added to the platform. In the near future, we plan to expand the usability of DECONbench by offering the possibility for owners of biological data to upload them. This extended functionality will allow health professionals and biologists to benefit from all developed methods to gain insights regarding the composition of their samples.

## Acknowledgements

We thank all members of the HADACA consortium for helpful discussion and contributions during the HADACA data challenge 2ndedition (November 2019, Aussois, France).

We also thank Daniel Jost and the members of the BCM team for inspiring discussions during regular joint group meetings. We are grateful to the Codalab data challenge open source platform. The authors gratefully acknowledge the EpiMed core facility for their support and assistance in this work. This work is part of the national program Cartes d'Identité des Tumeurs supported by the Ligue Nationale Contre le Cancer. Where authors are identified as personnel of the International Agency for Research on Cancer / World Health Organization, the authors alone are responsible for the views expressed in this article and they do not necessarily represent the decisions, policy or views of the International Agency for Research on Cancer / World Health Organization.

## HADACA (Health Data Challenge) Consortium

Nicolas Alcala^6^, Alexis Arnaud^2^, Francisco Avila Cobos^7^, Luciana Batista^8^, Anne-Françoise Batto^9^, Yuna Blum^3^, Florent Chuffart^10^, Jérôme Cros^5^, Clémentine Decamps^1^, Lara Dirian^11^, Daria Doncevic^12^, Ghislain Durif^13^, Silvia Yahel Bahena Hernandez^14^, Milan Jakobi^10^, Rémy Jardillier^15^, Marine Jeanmougin^16^, Paulina Jedynak^10^, Basile Jumentier^1^, Aliaksandra Kakoichankava^17^, Maria Kondili^18^, Jing Liu^19^, Tiago Maie^20^, Jules Marécaille^11^, Jane Merlevede^21^, Maxime Meylan^322^, Petr Nazarov^23^, Kapil Newar^1^, Karl Nyrén^14^, Florent Petitprez^3^, Claudio Novella Rausell^14^, Magali Richard^1^, Michael Scherer^24^, Nicolas Sompairac^21^, Katharina Waury^14^, Ting Xie^25^, Markella-Achilleia Zacharouli^14^

**Affiliations:**

^1^Laboratory TIMC-IMAG, UMR 5525, Univ. Grenoble Alpes, CNRS, Grenoble, France

^2^Data Institute, Univ. Grenoble Alpes, Grenoble,France

^3^Programme Cartes d’Identité des Tumeurs (CIT), Ligue Nationale Contre le Cancer, Paris, France

^4^INSERM U1068 CRCM, Marseille, France

^6^Section of Genetics, International Agency for Research on Cancer (IARC-WHO), Lyon, France

^7^Center for Medical Genetics Ghent, Department of Biomolecular Medicine, Ghent University, Ghent, Belgium

^8^Innate Pharma, Marseille, France

^9^Equipe Cancer et Immunité- INSERM Centre de Recherche des Cordeliers, Paris, France

^10^Institute for Advanced Biosciences, CNRS UMR 5309, Inserm, U1209, Univ. Grenoble Alpes, F-38700 Grenoble, France

^11^Verteego, Paris, France

^12^Health Data Science Unit, BioQuant Center and Medical Faculty Heidelberg, Germany

^13^Université de Montpellier, CNRS, IMAG UMR 5149, Montpellier, France

^14^Uppsala University, SE-751 05, Uppsala, Sweden

^15^University Grenoble Alpes, CEA, INSERM, IRIG, Biology of Cancer Infection UMR_S 1036, 38000 Grenoble, France & University Grenoble Alpes, CNRS, Grenoble INP, GIPSA-lab, Institute of Engineering University Grenoble Alpes, 38000 Grenoble, France

^16^Department of Molecular Oncology, Institute for Cancer Research, Oslo University Hospital, The Norwegian Radium Hospital - Oslo, Norway

^17^Vitebsk State Medical University & NatiVita, Vitebsk, Belarus

^18^Centre de Recherche de St. Antoine, Paris, AP-HP

^19^Institut Curie, PSL Research University, Sorbonne Universités, UPMC Université Paris 06, CNRS, UMR144, Equipe Labellisée Ligue contre le Cancer, 75005 Paris, France

^20^Institute for Computational Genomics, Joint Research Center for Computational Biomedicine, RWTH Aachen University Medical School, Aachen, Germany

^21^Institut Curie, PSL Research University, Mines Paris Tech, Inserm, U900, F−75005, Paris, France

^22^INSERM U1138 Centre de Recherche des Cordeliers, France

^23^Quantitative Biology Unit, Luxembourg Institute of Health, L-1445 Strassen, Luxembourg

^14^Uppsala University, SE-751 05, Uppsala, Sweden

^24^Department of Genetics/Epigenetics, Saarland University, Saarbrücken, Germany

^25^Centre de Recherche en Cancérologie de Toulouse, Inserm UMR 1037, F-31037, Toulouse, France

