## SUPPLEMENTARY INFORMATION for "DECONbench: a benchmarking platform dedicated to deconvolution methods for tumor heterogeneity quantification"

### Reference benchmarking methods:

Eight methods have been included in the benchmarking platform as reference methods to be compared with the accuracy of new methods.

The method RNA_wICA (r_WIC) relies on transcriptomic data and uses Independent Component Analysis (ICA) for both feature selection and deconvolution. It relies on the use of the functions “runICA” and “getGenesICA” by P. Nazarov (sablab.net/scripts/LibICA.r) and the deconica R package. For the ICA-based feature selection, the function “runICA” is run for the first time with the parameters ncomp = 10 and ntry = 50. Then, the function “getGenesICA” selects top-contributing genes with a FDR of 0.2, the feature selection is done on these contributing genes belonging to a component having an average stability greater than 0.8. Finally, duplicated genes are removed.
For the ICA-based deconvolution step, the function deconica::run_fastica is run with parameters overdecompose = FALSE and n.comp = 5, remaining parameters set to default. The 30 most important genes of each ICA component are extracted by the function deconica::generate_markers with the parameter return = "gene.ranked". These genes are used to weight the components score with the function deconica::get_scores, with the log counts of the ICA as “df” parameter, the list of 30 genes as “markers.list” parameter, and the parameter summary = "weighted.mean". Finally, the proportions are extracted with the function deconica::stacked_proportions_plot on the transpose.

The method RNA_wNMF (r_WNM), using transcriptomic data, is in two steps. The first step uses ICA to perform a feature selection as described for RNA_wICA, although duplicated genes are kept. This step, therefore, allows a gene which contributes to several components to be present several times in the data. The deconvolution is based on Non-negative Matrix Factorization (NMF) called by the NMF::nmf function, with the parameter method = "snmf/r".

The method DNAm_wICA (m_WIC) analyzes DNA methylation data. It does not provide feature selection. The deconvolution step is based on ICA, similarly to what was described for the second step of RNA_wICA, but applied on the DNA methylation matrix.

The method DNAm_EDec (m_EDC), which estimates sample composition from DNA methylation data, follows the pipeline implemented in the R package medepir (HADACA consortium et al., 2020). The feature selection is made with medepir::feature_selection to keep the probes with a variance higher than 0.02. The method EDec (Onuchic et al., 2016) is used for the deconvolution part, with the function medepir::Edec and all the selected probes as “infloci” parameter.

The method DNAm_MeDeCom (m_MDC), also using DNA methylation, is similarly based on the pipeline of the R package medepir. The feature selection is made as for DNAm_EDec above, on the probes with a variance higher than 0.02. The deconvolution step, however, is from the MeDeCom R package (Lutsik et al., 2017). It is run with the function MeDeCom::runMeDeCom, with the lambda parameter set to 0.01.

The method both_wICA (b_WIC) combines the input of both transcriptomics and DNA methylation. It has no feature selection step. The deconvolution is in two steps, one on each data type. The transcriptomics and DNA methylation are separately deconvoluted with the same deconvolution step as in r_WIC and m_WIC respectively. Finally, the mean of both deconvolution matrices is computed as the final method output. To compute the average, the cell types of the both deconvolution matrices are matched by iteration. The cell types of the methylation result matrix are reordered 1,000 times, and the one that best correlates with the transcriptomic result matrix is kept.

The method both_wNMFMeDeCom (b_COM), also combining transcriptomics and DNA methylation, is the combination of the two methods RNA_wNMF and DNAm_MeDeCom. The method r_WNM is first applied on the RNAseq matrix. The DNA methylation matrix is pre-treated as described in the m_MDC method, with the selection of probes with a variance higher than 0.02. Finally, the method MeDeCom is run on the DNAm data, with the result of r_WNM as initialization parameter startA.

Finally, the method both_meanwNMFMeDeCom (b_MEA), which integrates transcriptomics and DNA methylation, applies r_WNM to the transcriptomics matrix, m_MDC to the DNA methylation matrix, and returns the mean of the two results matrices, similarly to what we explained for b_WIC.

**Supplementary Table 1**: Description of each reference benchmarking method.

| **Name** | **RNA_wICA** | **RNA_wNMF** | **DNAm_EDec** | **DNAm_MeDeCom** | **DNAm_wICA** | **both_wICA** | **both_wNMFMeDeCom** | **both_meanwNMFMeDeCom** |
|---|---|---|---|---|---|---|---|---|
| **Acronym** | **r_WIC** | **r_WNM** | **m_EDC** | **m_MDC** | **m_WIC** | **b_WIC** | **b_COM** | **b_MEA** |
| **Data type** | RNA | RNA | DNAm | DNAm | DNAm | both | both | both |
| **FS DNAm** | / | / | Var > 0.02 | Var > 0.02 | / | / | Var > 0.02 | Var > 0.02 |
| **FS RNA** | ICA, most important genes of most stable components, removing of duplicated genes | ICA, most important genes of most stable components, not removing of duplicated genes | / | / | / | / | ICA, most important genes of most stable components, not removing of duplicated genes | ICA, most important genes of most stable components, not removing of duplicated genes |
| **Deconvolution DNAm** | / | / | Edec | MeDeCom | ICA weighted on 30 most important genes | ICA weighted on 30 most important genes | MeDeCom with the A matrix computed on RNA as startA parameter | MeDeCom |
| **Deconvolution RNA** | ICA weighted on 30 most important genes | NMF with snmf/r method | / | / | / | ICA weighted on 30 most important genes | NMF with snmf/r method | NMF with snmf/r method |
| **Time 10 A** | ~10mn | ~20mn | ~3h | ~17h | ~10mn | ~10mn | ~17h | ~17h30 |
| **Time 1 A** | ~1mn | ~2mn | ~20mn | ~1h40 | ~1mn | ~1mn | ~1h40 | ~1h45 |

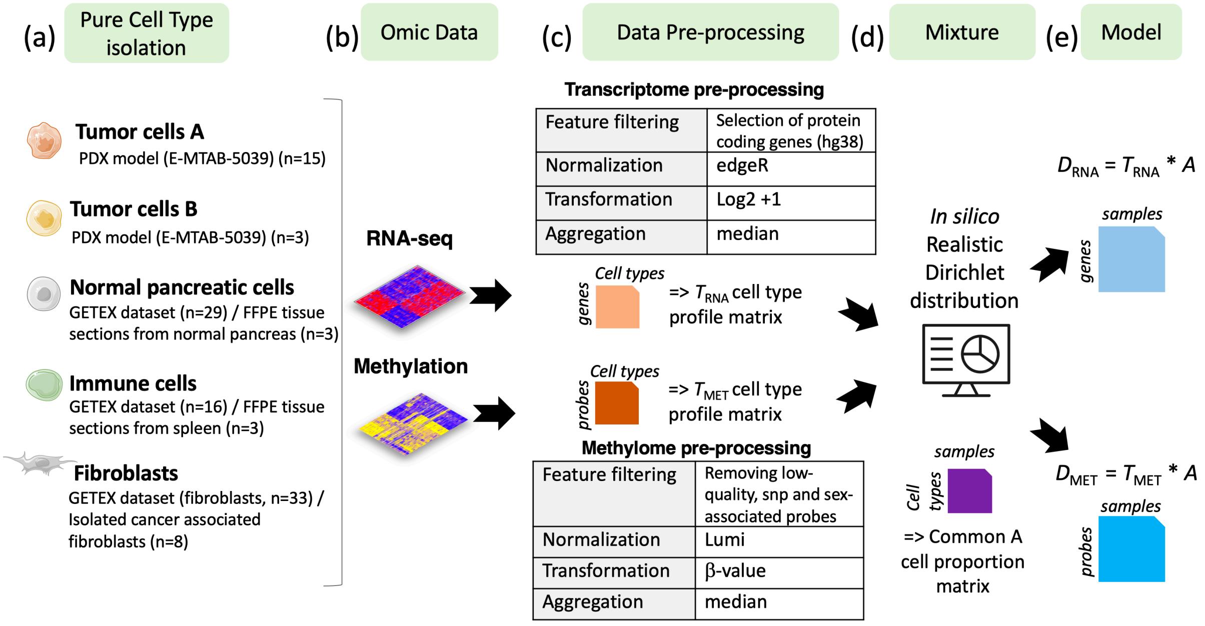


**Supplementary Figure 1. Benchmark dataset construction**: (a) 5 different cell populations present in pancreatic tumors were considered. (b) Raw transcriptome and methylome profiles of these different cell populations were extracted from various sources (PDX model, tissues or isolated cells). (c) Raw cell type profile matrices were preprocessed (Feature filtering, normalization, signal transformation, sample aggregation) to avoid any batch effect. After pre-processing, transcriptomic data are constituted of log2-transformed expression counts on 21,566 genes and methylome data of beta-values on 772,316 EPIC array CpG sites. (d) In silico Dirichlet distribution have been used based on realistic proportions defined by the anatomopathologist expertise (J. Cros). (e) Paired methylome and transcriptome of in silico mixtures from pancreatic tumors were obtained by considering $D = T A$, with T the cell-type profiles (matrix of size M * K, with M the number of features and K=5 the number of cell types) and A the cell-type proportion per patient (matrix of size K * N, with N=30 the number of samples) common between both omics. One training set ($D_{MET}$ *and* $D_{RNA}$) is accessible to the users (obtained by one realization of A). The algorithms are compared on 10 test sets (obtained from 10 other realizations of A) that are hidden on the platform.

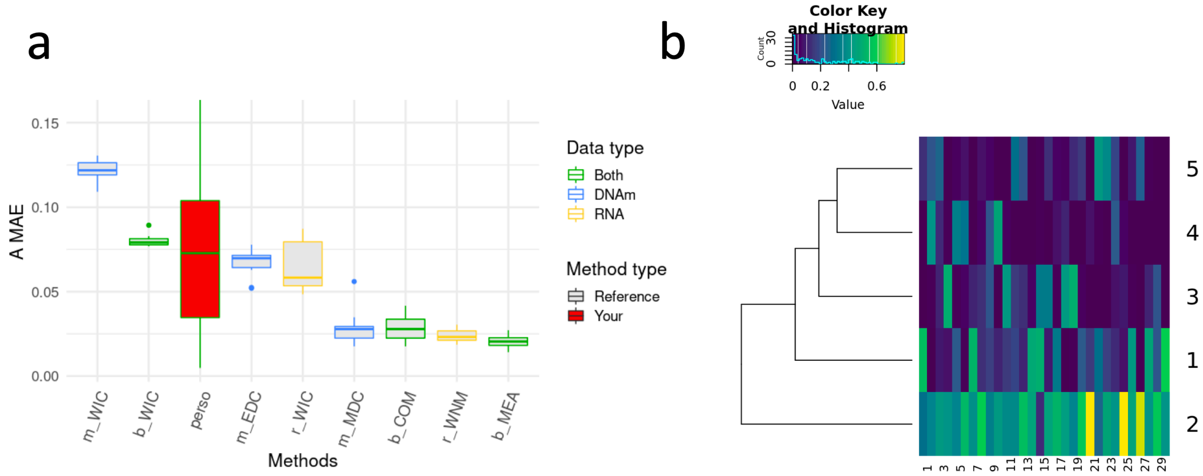


**Supplementary Figure 2. DECONBench graphical outputs**. (a) Boxplots of the Mean Absolute Errors (MAE) of the estimations of the A matrices (i.e. proportion matrices) obtained for each method that uses the transcriptome only (yellow), the methylome only (blue) or both omics (green). Boxplots of the reference methods and other existing methods are shown in grey, whereas the user's method is shown in red. (b) Heatmaps of each A-matrix estimate are generated. The cell populations are in rows and the samples in columns.
